# Chemical degradation identified Human p38-MAPK as a host target to combat parasitic infections

**DOI:** 10.1101/2024.12.19.629355

**Authors:** Jhalak Singhal, Neha Rawat, Ruby Bansal, Pallavi Srivastava, Gajala Deethamvali Ghouse Peer, Ramendra Pati Pandey, Anand Ranganathan, Shailja Singh

## Abstract

The interplay between host and parasite determines parasite burden and disease outcome. Parasite exploits host signaling pathways like p38-MAPK for its survival and pathogenesis. NR-7h, a proteolysis-targeting chimera (PROTAC) targeting human p38-MAPK was used to assess p38-MAPK’s role in *Leishmania donovani* and *Plasmodium falciparum* infection in their respective hosts. NR-7h degraded host p38-MAPK in a time- and dose-dependent manner. Degradation of host p38-MAPK by NR-7h reduced parasite load in host cells dose-dependently, implicating the role of p38-MAPK in parasite survival. The modulation of cytokine profiling and oxidative burst upon NR-7h mediated degradation of host p38-MAPK was further correlated with parasite death. The synergistic effect of host p38-MAPK degradation by NR-7h with Amphotericin B enhances the efficacy of parasite-directed therapy. This study underscores the importance of host p38-MAPK for *L. donovani* and *P. falciparum* progression and highlights NR-7h’s potential in antiparasitic therapy by targeting this pathway.

## Introduction

*Leishmania donovani (L. donovani)*, a protozoan causes Visceral Leishmaniasis (VL) that has been classified as a threat to human beings causing loss of weight and fever with spleen and liver swelling^1,2^. Annually, around 50,000 to 90,000 new cases of VL are reported worldwide^3^. The disease outcome always depends upon the early response from the host cells to encounter the pathogen including the production of pro-inflammatory responses^4,5^, generation of oxidative bursts^6,7^, and phagocytosis^8,9^. However, it is well reported that *Leishmania* parasites evade host immune responses by modulating host factors/signaling that includes more production of anti-inflammatory cytokines^10,11^, induction of autophagy^12–15^, and hijacking of post-translational modifications like Sumoylation^16^. *Plasmodium falciparum (P. falciparum)* parasitizes human erythrocytes and causes malaria which is a global health burden with an estimated 249 million cases and about 6 million deaths in 2022^17^. The complex life cycle, high polymorphism, immune evasion, and host modulation are some of the challenges that impede the Malaria elimination program. Many efforts have been made to develop therapeutics targeting parasites, however, due to the development of drug resistance, these therapeutics are not effective^18,19,20^.To overcome this issue, there is an urgent need to investigate host targets to develop effective therapeutics.

The human p38-MAPK pathway plays a critical role in various cellular processes such as inflammation, cell growth, differentiation, and cell death^21,22^. Upon activation, p38-MAPK modulates pro-inflammatory cytokines, chemokines, and adhesion molecules that are responsible for recruiting the immune cells at the site of infection^23^. Also, p38-MAPK modulates the expression of the antimicrobial peptides to improve the phagocytosis process of immune cells and thus a better presentation of antigens to major histocompatibility complexes^24^. Moreover, p38-MAPK has served as an adjunctive therapy for the management of inflammatory disorders and immune responses induced to combat infections^25,26^. There are reports on the inhibition or modulation of p38-MAPK activity as a potential enhancing agent for an appropriate immune response^27^. At the cellular level, host p38-MAPK signaling is known to be modulated during parasite infections which further emphasizes the need to study the disease pathogenesis in absence of host p38-MAPK. In context of Leishmaniasis, it is reported that human p38-MAPK also plays a role in inflammation and immunomodulation, thus regulating the progression and severity of infection of the *Leishmania* parasites^28,29^. Studies have shown that p38-MAPK activity modulates the intracellular survival and multiplication of *Leishmania* parasites in human macrophages^30–33^.

*Plasmodium falciparum* infection in human erythrocytes also involves constant interaction between host and parasite. Multiple studies have shown that during the asexual stage, the parasite modulates host signaling pathways for the disease progression^34–36^ highlighting the targets for host-directed therapies. The parasite exports its protein into the erythrocytes resulting in remodelling of the erythrocyte and altercation in the architecture of the erythrocyte membrane^37^. By doing so, the parasite exploits the host cell for its own growth and survival including nutrient uptake^38^, antigen presentation^39^, and interaction with cytoskeletal proteins^40^, affecting the normal erythrocyte physiology. These modulations help the parasite to evade the immune system, increase its virulence, and support its survival within the host cell. Mature human erythrocytes lack nuclei and organelles and thus fail to maintain homeostasis by genetic regulation. In normal erythrocyte physiology, p38-MAPK maintains membrane stability and responds to oxidative or osmotic stress. Under hyperosmotic stress, p38-MAPK activation results in induction of eryptosis and its inhibition by p38-MAPK inhibitors results in the rescuing of erythrocytes from classical hallmarks of eryptosis including phosphatidylserine exposure, intracellular calcium levels and ROS^41^. Its activation is tightly regulated and involves phosphorylation by upstream kinases including MAP kinase kinase (MKK3/6)^42^ in response to stress signals, leading to downstream effects such as phosphorylation of cytoskeletal and membrane proteins. Under pathological conditions like malaria, erythrocyte p38-MAPK assumes a more dynamic role. Therefore, targeting human p38-MAPK could be a potential therapeutic strategy to impede the disease progression caused by the sister parasites *Leishmania* and *Plasmodium*.

Due to the selectivity and high toxicity, many p38-MAPK inhibitors have been unsuccessful at the clinical trial stage^43^. Recently, proteolysis-targeting chimeras or PROTACs have emerged as a new therapeutic strategy for targeted protein degradation owing to their high specificity. A PROTAC is an innovative therapeutic modality that takes advantage of bifunctional molecules selectively targeting proteins for degradation. PROTACs recruit an E3 ubiquitin ligase to bind with the target protein; it ubiquitinates and degrades it through the proteasome^44,45^. PROTACs fundamentally differ from traditional inhibitors in that, while a conventional inhibitor can inhibit the protein’s function, PROTACs degrade the protein inside the cell. This can, therefore, allow disease treatment by degrading targeted proteins rather than suppressing their activity. Moreover, PROTACs are beneficial to target proteins conventionally considered “undruggable”^46^.

Using the PROTAC technology, we have explored a one-of-its-kind inhibitor that targets and degrades p38-MAPK^47^, which plays a vital role in the invasion and survival of the *L. donovani* and *P. falciparum* in their respective hosts. This approach might be more effective with specificity compared to present modes of treatment for visceral leishmaniasis and malaria as it reduces the risk associated with potential drug resistance and toxicity in side effects.

## Results

### Degradation of human macrophage p38-MAPK via PROTAC molecule, NR-7h

To investigate the activation levels of human macrophage p38-MAPK upon *L. donovani* infection in macrophages, we infected THP-1 macrophages with 20 MOI of *L. donovani* promastigotes over varying time intervals. The expression level of phospho-p38-MAPK (P-p38-MAPK) was monitored by western blotting at different time points. The data shows no significant alterations in the expression level of P-p38-MAPK in the infected macrophages compared to the uninfected macrophages at 2hr and 4hr post-infection. However, a significant upregulation (∼1.5-fold) in the P-p38-MAPK expression was observed in infected macrophages at 6hr post-infection. Interestingly, after 12hr of infection, the expression level of P-p38-MAPK was found to be significantly downregulated (∼1.5-fold) in the infected macrophages compared to uninfected macrophages (Fig. 1A), suggesting the activation of human macrophage p38-MAPK during the early stages of *L. donovani* invasion.

**Fig. 1:**
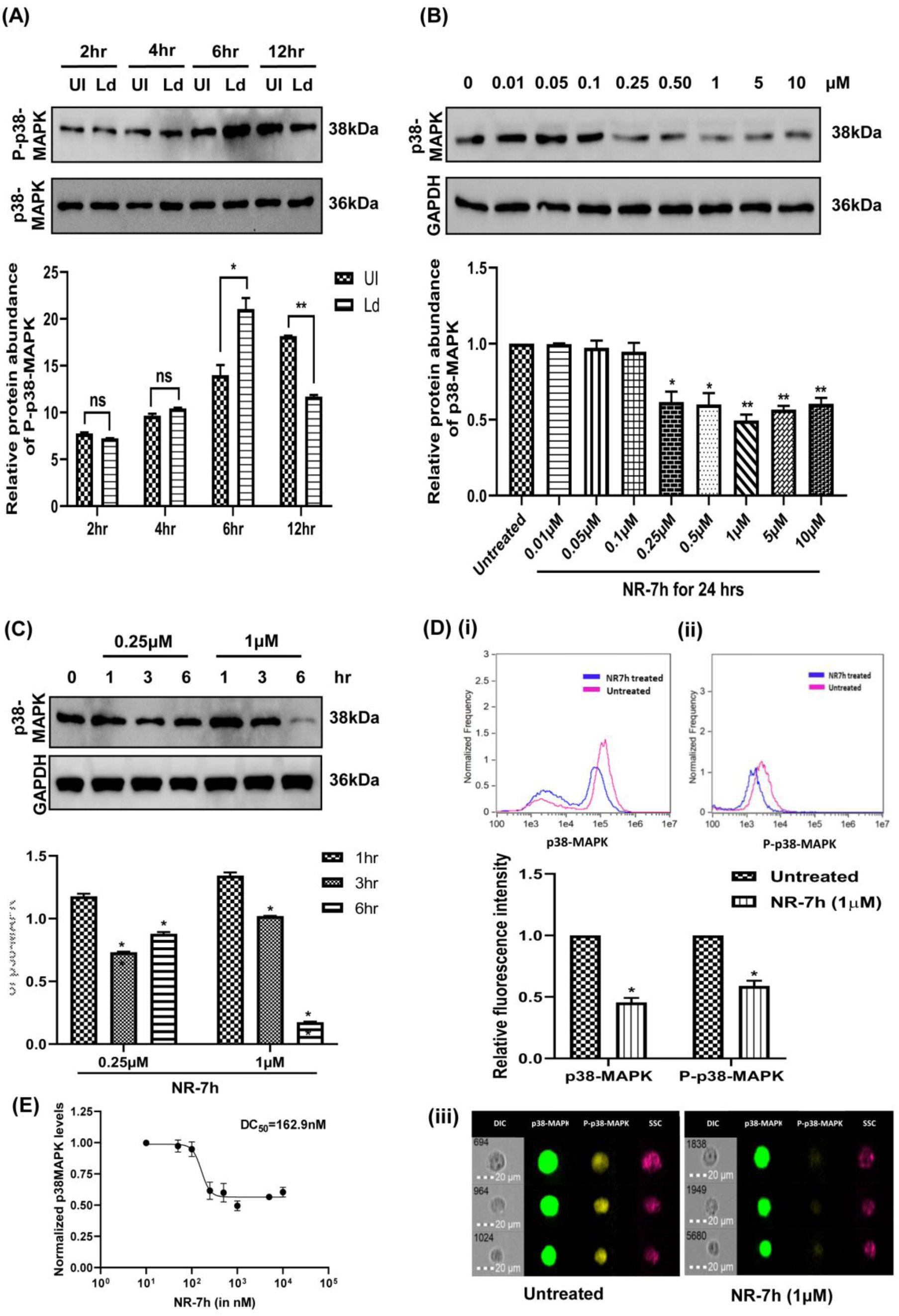
(A) Expression of p38-MAPK in infected macrophages. PMA-differentiated THP-1 macrophages were infected with 20 MOI of *L. donovani* for varying durations (2hr, 4hr, 6hr, and 12hr). Cell lysates were prepared from the infected macrophages. Western blotting was employed using a specific antibody to quantify the expression of p38-MAPK. **(B-D) Degradation of p38-MAPK by NR-7h** PMA differentiated THP-1 macrophages were treated with different concentrations of NR-7h concurrently with the stimulation of 100ng/mL of LPS for 24hr **(B)** and 1hr, 3hr, and 6hr **(C)**. Western blotting was used to evaluate the levels of p38-MAPK expression. Untreated cells were used as control. **(D)** Imaging flow cytometry was used to evaluate the fluorescence levels of **(i)** p38-MAPK, **(ii)** P-p38-MAPK **(iii)** Image panel of macrophages. **(B-C)** GAPDH served as the loading control. Data from one of three experiments were presented. Band intensities were analyzed using ImageJ software and plotted using GraphPad Prism 8. Statistical analysis utilized the unpaired t-test with Welch’s correction, with significance levels denoted as *, **, and *** for p-values less than 0.05, 0.01, and 0.001, respectively. **(E)** The half-maximal degradation concentration (DC_50_) of NR-7h was calculated and plotted in GraphPad Prism 8.

To study the functional role of human macrophage p38-MAPK during *L. donovani* infection, we applied an advanced approach of utilizing the PROTAC molecule, NR-7h. THP-1 macrophages were treated with gradient concentrations of NR-7h for 24hr and to check the degradation ability, p38-MAPK protein levels were monitored by western blotting. Significant degradation of p38-MAPK (∼2-fold) was observed upon the treatment of NR-7h across concentrations ranging from 0.25μM to 10μM at 24hr post-treatment (Fig. 1B). Furthermore, we evaluated the degradation of human macrophage p38-MAPK upon the treatment of NR-7h at shorter time points (Fig. 1C). We found that upon treatment of NR-7h at the concentrations of 0.25μM and 1μM for 1hr, 3hr, and 6hr in the macrophages, the maximal degradation of p38-MAPK was observed upon treatment of 1μM of NR-7h for 6hr in a concentration- and time-dependent manner. We also observed a lower expression of p38-MAPK and P-p38-MAPK in 1μM NR-7h-treated macrophages compared to untreated macrophages using imaging flow cytometry (Fig. 1D). The half-maximal degradation concentration (DC50) of NR-7h was noted to be 162.9nM in macrophages (Fig. 1E).

### NR-7h-mediated degradation of human macrophage p38-MAPK reduced the parasitic burden of *L. donovani*

To check the cytotoxic effect of NR-7h on macrophage viability, we performed an MTT assay and found no significant alterations in the viability of macrophages upon treatment with various concentrations of NR-7h (Fig. 2A). As depicted in Fig. 1A, the expression level of P-p38-MAPK was upregulated at 6hr post-*L. donovani* infection, therefore we further investigated the role of human macrophage p38-MAPK in the invasion of *L. donovani*. Different concentrations of NR-7h were employed to degrade human macrophage p38-MAPK for 6hr followed by the infection of *L. donovani* parasites. The intracellular parasite load was calculated by quantifying the relative expression level of the kinetoplastid minicircle housekeeping gene (JW gene) of *L. donovani* using quantitative Real-Time PCR (qRT-PCR). The maximum clearance of the intracellular parasites was observed at the concentration of 0.75μM (∼16-fold), and 1μM (∼8-fold) of NR-7h in infected macrophages. (Fig. 2B). Confocal microscopy was also used to count the number of intracellular amastigotes and to calculate the infectivity rate in NR-7h-treated-infected macrophages as compared to the untreated-infected macrophages (Fig. 2C(i)). A significant reduction of *L. donovani* infected cells was observed upon the treatment of NR-7h on the concentrations of 0.5μM and 1μM (Fig. 2C(ii)). Similarly, a significant reduction in intracellular amastigotes was observed in a dose-dependent treatment of NR-7h (Fig. 2C(iii)). These results underscore the pivotal role of human macrophage p38-MAPK in favoring *L. donovani* invasion dynamics.

**Fig. 2:**
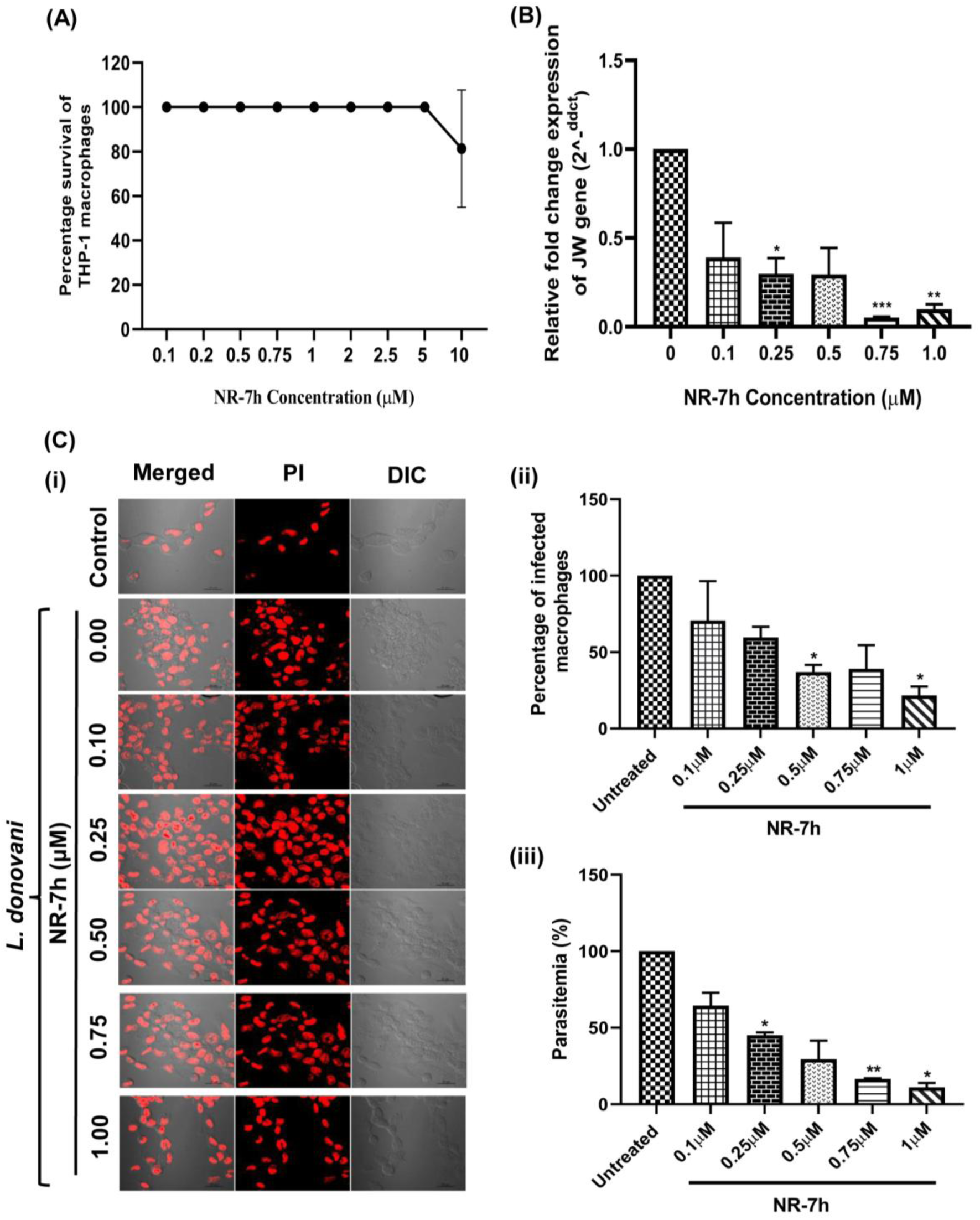
Effect of NR-7h on the parasite burden in infected macrophages. **(A)** PMA differentiated THP-1 macrophages were treated with varying concentrations of NR-7h for 72hr. The viability of cells was assessed using the MTT assay. **(B and C)** PMA-differentiated THP-1 macrophages were treated with different concentrations of NR-7h in conjunction with the stimulation of 100 ng/ml LPS for 6hr, followed by infection with 20 MOI of *L. donovani* for 24hr. Untreated infected macrophages served as the control group. **(B)** RNA was isolated to quantify the expression of the JW gene using qRT-PCR. **(C) (i)** Parasite load was measured by using Propidium iodide (PI) staining via confocal microscopy. Scale bar, 20µm. **(ii)** Percentage of infected macrophages and **(iii)** Percent parasitemia were plotted using GraphPad Prism 8. Data represents the mean ± standard deviation (SD) from three independent experiments. Statistical significance was determined using Student’s unpaired 2-tailed t-test, with significance levels denoted as *, **, and *** for p-values less than 0.05, 0.01, and 0.001, respectively.

Next, we investigated the effect of NR-7h treatment on the ability of human macrophage p-38 MAPK degradation and subsequently on the parasite load till 60hr of *L. donovani* infection. We observed the significant degradation of human macrophage p-38 MAPK in terms of less fluorescence intensity at 12hr, 36hr, and 60hr post-infection. Also, the reduction in the intracellular parasite load was observed upon NR-7h treatment till 60hr of infection. The data shows the efficacy of NR-7h treatment on p38-MAPK degradation and subsequent reduction of the parasite load in a time-dependent manner (Figure 3).

**Fig. 3:**
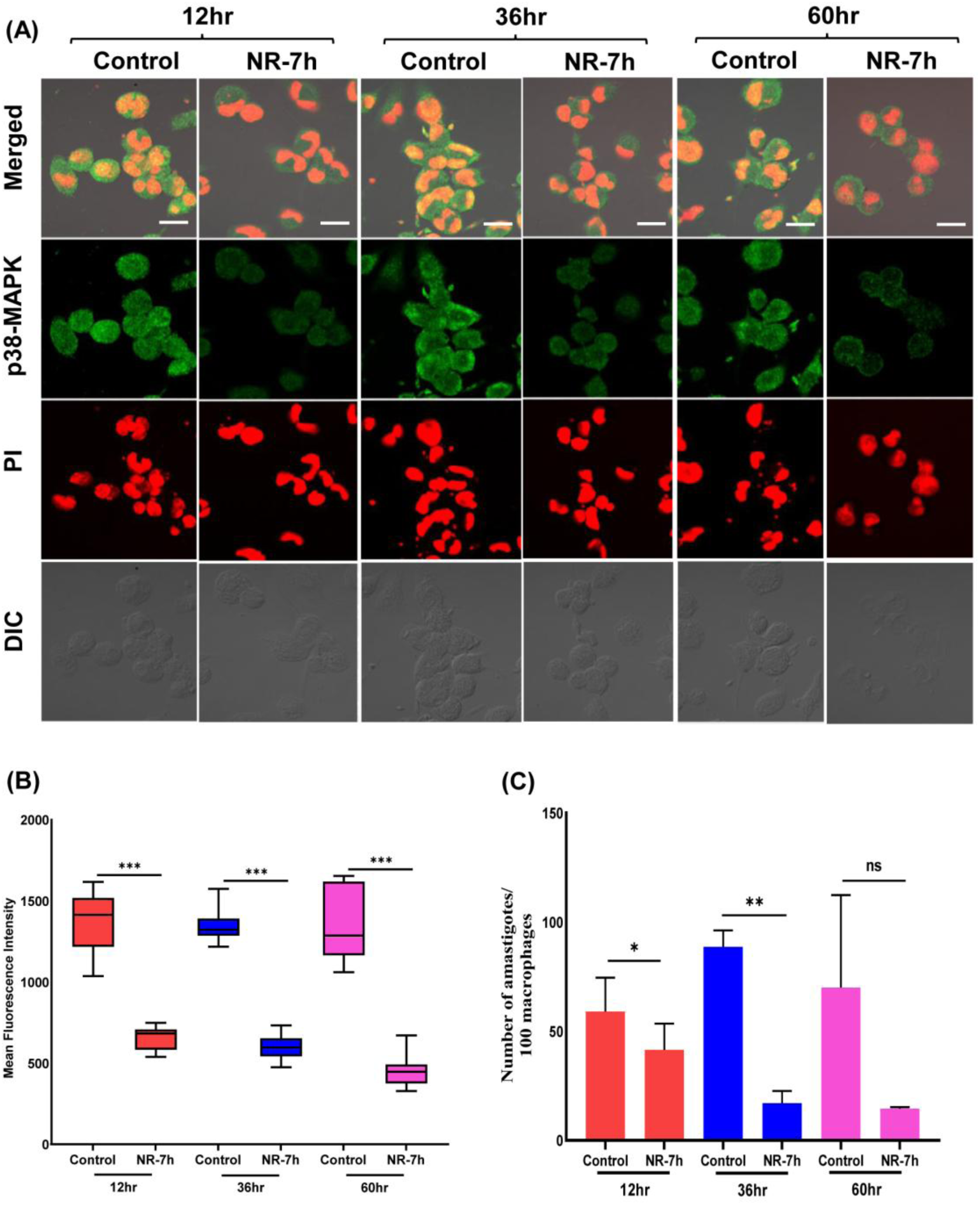
Long-term effect of NR-7h treatment on p38-MAPK degradation as well as on parasite burden. **(A)** PMA-differentiated THP-1 macrophages were treated with 1μM of NR-7h in conjunction with the stimulation of 100 ng/mL LPS for 6hr, followed by infection with 20 MOI of *L. donovani* for 12hr, 36hr, and 60hr. Untreated infected macrophages served as the control group. The green fluorescence intensity of p38-MAPK by staining with Alexa fluor 488 while parasite load by staining with propidium iodide (PI) was monitored by confocal microscopy. Scale bar, 20µm. Data from one of three experiments were presented. **(B)** The mean fluorescence intensity of p38-MAPK and **(C)** parasite burden was plotted using GraphPad Prism 8. Data represents the mean ± standard deviation (SD) from three independent experiments. Statistical significance was determined using Student’s unpaired 2-tailed t-test, with significance levels denoted as *, **, and *** for p-values less than 0.05, 0.01, and 0.001, respectively.

### Degradation of human macrophage p38-MAPK by NR-7h modulates host immune mechanisms during *L. donovani* infection

The p38-MAPK pathway is an essential modulator of numerous cellular signaling processes, including inflammation and cytokine production^48,49^. Here, we explored how the degradation of human p38-MAPK alters cytokine secretion and presents complex dynamics of regulation. Treatment of human macrophages with different doses of NR-7h drastically suppressed the production of TNFα, a proinflammatory cytokine as well as the anti-inflammatory cytokine, IL-10 compared to the untreated control. Interestingly, when *L. donovani* was introduced to NR-7h-pretreated macrophages, a higher upregulation in the production of TNFα (∼4-fold) was observed compared to the untreated infected macrophages. However, a significant decrease in the production of IL-10 (∼4-fold) was also observed compared to the untreated infected macrophages (Fig. 4A). These findings reveal the interplay between human p38-MAPK activity and cytokine profile that regulates the infection of *L. donovani*, thereby suggesting the pathogen’s adaptive mechanisms for developing susceptibility and pathogenicity in macrophages.

**Fig. 4:**
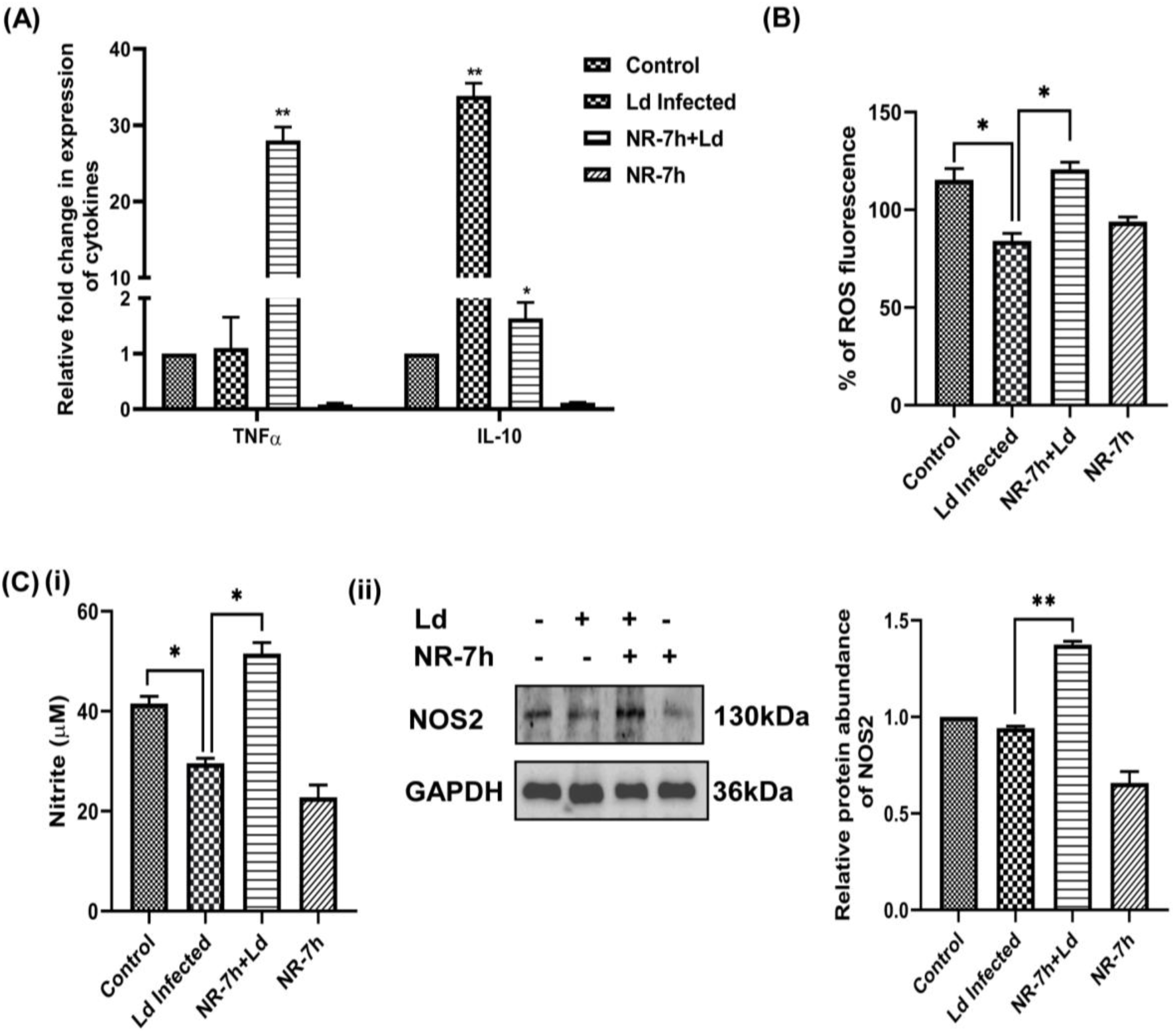
Effect of NR-7h treatment on the immune mechanisms in infected macrophages. **(A)** PMA-differentiated THP-1 macrophages were treated with 1μM of NR-7h, alongside stimulation with 100 ng/mL LPS for 6hr followed by infection with 20 MOI of *L. donovani* for 24hr. RNA was isolated to quantify the expression level of inflammatory markers using qRT-PCR. Untreated macrophages served as the control group. Data represents the mean ± standard deviation (SD) from three independent experiments. Statistical significance was determined using Student’s unpaired 2-tailed t-test. **(B)** PMA-differentiated THP-1 macrophages were treated with 1μM of NR-7h, along with stimulation with 100 ng/mL LPS for 6hr, followed by infection with *L. donovani* for 30 min, with the addition of 10μM DCFDA. Samples were analyzed using fluorimetry. **(C)** PMA-differentiated THP-1 macrophages were treated with 1μM of NR-7h, in combination with 100 ng/mL LPS and 20 ng/mL of hIFNγ for 6hr, followed by infection with *L. donovani* for 24hr. (i) Nitrite levels were measured using the supernatant with Griess reagent. (ii) Cell lysates were prepared to assess the expression level of NOS2, with GAPDH utilized as the loading control. Data presented is from one of three experiments. Band intensities were quantified using ImageJ software and plotted using GraphPad Prism 8. Statistical significance was assessed using the unpaired t-test with Welch’s correction, with significance levels denoted as *, **, and *** for p-values less than 0.05, 0.01, and 0.001, respectively.

Oxidative burst-mediated responses of human macrophages are critical regulatory events determining *L. donovani* survival and infectivity, and the generation level of reactive oxygen species (ROS) is closely linked with p38-MAPK activation. Using fluorimetry, we observed a significant downregulation of ROS generation in human macrophages when infected with *L. donovani*, however a spectacular upregulation of ROS generation amounts when the macrophages were pretreated with NR-7h for 6hr, followed by the infection of *Leishmania* parasites for 30 mins (Fig. 4B). Consequently, Nitric Oxide (NO) production was observed wherein a significant reduction in nitrite levels was noticed in human macrophages post 24hr of -*L. donovani* infection, however, NR-7h-treated-*L. donovani* infected macrophages showed a significant increase in nitrite levels (Fig. 4C(i)). Since Nitric Oxide Synthase 2 (NOS2) catalyzes the production of NO, its expression levels were also monitored and observed to be upregulated (∼1.5-fold) in NR-7h treated-infected macrophages (Fig. 4C (ii)), compared to untreated-infected macrophages.

Altogether, the data obtained provides new insights into complex interactions between *L. donovani* and the human macrophage p38-MAPK-mediated immune response during invasion, thus opening new perspectives in the parasite-human macrophage interaction and therapeutic strategies.

### Combinatorial efficacy of NR-7h and Amphotericin B against intra-macrophagic amastigotes in *L. donovani*-infected macrophages

Our study demonstrates the anti-leishmanial effect of p38-MAPK degradation by NR-7h in *L. donovani-*infected macrophages. Next, we investigated the anti-leishmanial ability of NR-7h in combination with Amphotericin B. Macrophages were pre-conditioned with diverse concentrations of NR-7h for 6hr before infection with *L. donovani* and concurrently treated with Amphotericin B at a concentration of 10nM for 24hr. qRT-PCR was used to quantify the intracellular parasite burden in measuring the relative expression levels of the kinetoplastid minicircle housekeeping gene (JW gene) of *Leishmania*. The parasite load was reduced by approximately 70%, in a dose-dependent manner, when infected cells were treated with either 10nM Amphotericin B alone or the different doses of NR-7h alone. Treatment of *L. donovani*-infected macrophages with NR-7h alone resulted in a dosage-dependent reduction of parasite burden. The combination of Amphotericin B (10nM) and NR-7h (0.5μM/1μM) significantly enhanced the clearance of intracellular parasites in a synergistic manner (Fig. 5A). These synergistic effects translated to a more significant decrease in parasite load than either modality alone and made the possibility of integrating human macrophage-directed strategies, such as p38-MAPK degradation via NR-7h with conventional anti-leishmanial therapeutics. No cytotoxicity effect was observed upon the treatment of both compounds in the macrophages (Fig. 5B).

**Fig. 5:**
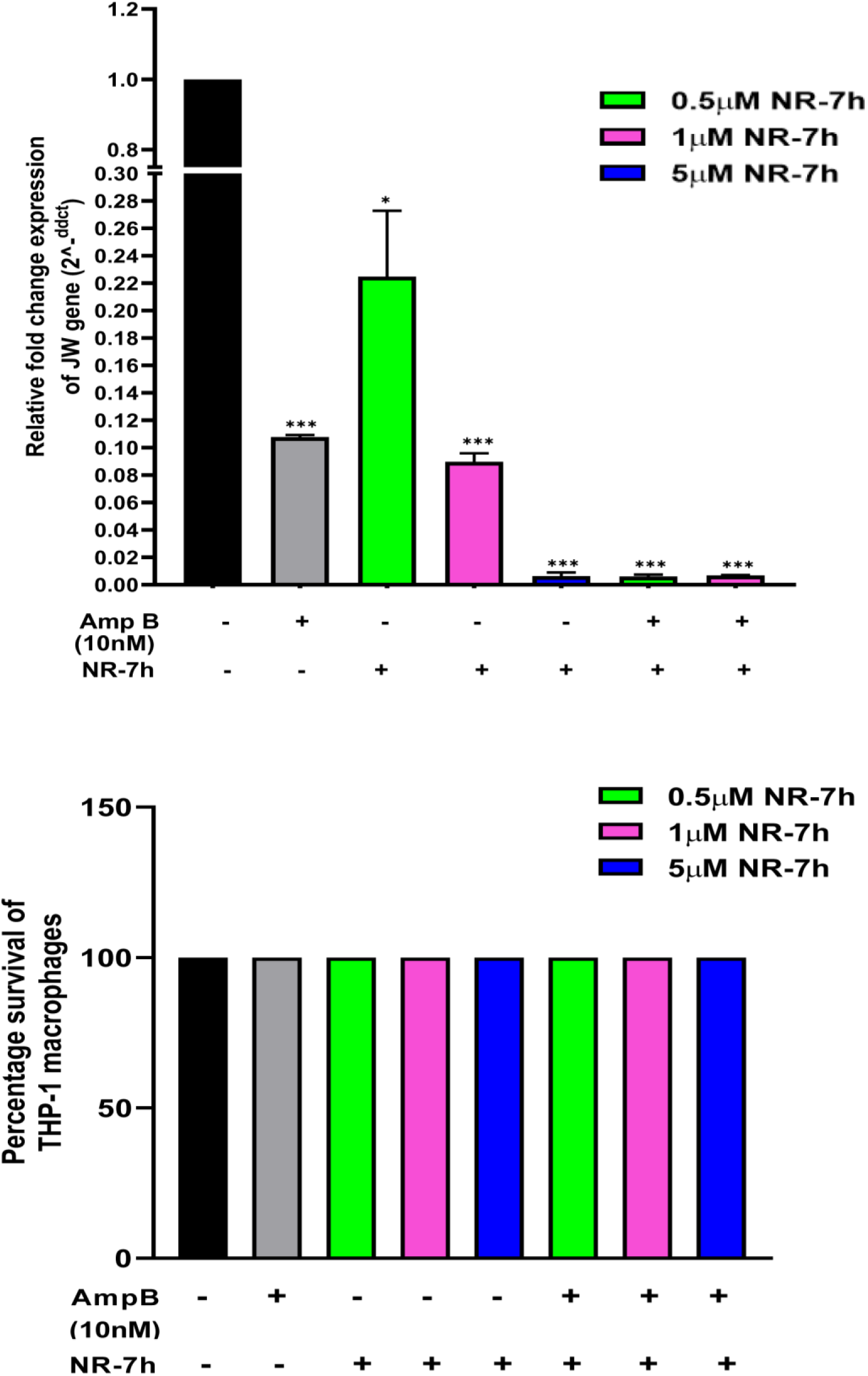
Effect of combination of NR-7h and Amphotericin B treatment in *L. donovani* infected macrophages. **(A)** PMA-differentiated THP-1 macrophages were treated with various concentrations of NR-7h along with the stimulation with 100 ng/mL LPS for 6hr, while another group received only 10nM Amphotericin B treatment for 24hr. For the combination treatment group, macrophages pre-treated with NR-7h were infected with *L. donovani* at 20 MOI and simultaneously treated with 10nM Amphotericin B. RNA was isolated to assess the expression level of the JW gene using qRT-PCR. **(B)** The viability of uninfected THP-1 macrophages treated by the combination of NR-7h and Amphotericin B was assessed using the MTT assay. Data presented is the mean ± standard deviation (SD) from three independent experiments. Statistical analysis was performed using Student’s unpaired 2-tailed t-test, with significance levels denoted as *, **, and *** for p-values less than 0.05, 0.01, and 0.001, respectively.

### NR-7h mediated degradation of p38-MAPK in human erythrocytes and its antimalarial potential

To investigate the effect of erythrocyte p38-MAPK degradation on *Plasmodium* infection, NR-7h PROTAC was deployed. The erythrocytes were treated with different concentrations of NR-7h (1µM, 2.5µM, and 5µM) and the expression of p38-MAPK was assessed by immunoblotting. The results suggested that NR-7h significantly degraded erythrocyte p38-MAPK (∼2-fold) compared to untreated control at all the concentrations (Fig. 6A (i)). Additionally, reduced expression of p38-MAPK in NR-7h (2.5µM) treated erythrocytes was confirmed by imaging flow cytometry (Fig. 6A (ii)). Further, to confirm the ubiquitin-proteasome-dependent mechanism of NR-7h, erythrocytes were treated with NR-7h (2.5µM) in the absence and presence of MG132, a proteasomal inhibitor. Immunoblotting revealed that the p38-MAPK degradation was significant (2-fold) in NR-7h-treated erythrocytes in the absence of MG132 (Fig. 6B). Additionally, the ability of NR-7h to degrade host erythrocyte p38-MAPK in *P. falciparum*-infected erythrocytes was also assessed by immunoblotting (Fig. 6C (i)).

**Fig. 6:**
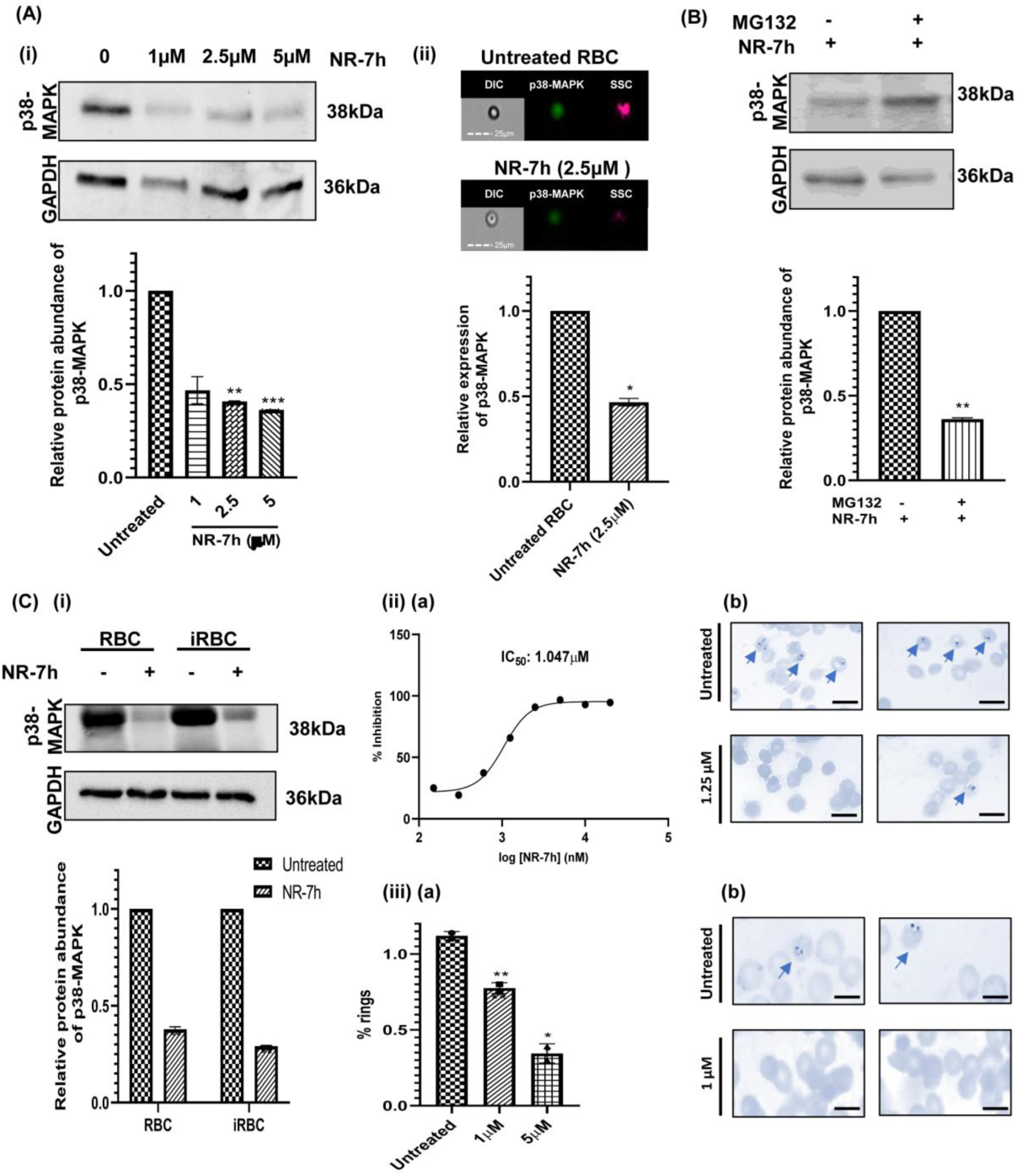
Degradation of erythrocyte p38-MAPK by NR-7h and its antimalarial potency. **(A)** **(i)** Uninfected erythrocytes were treated with different concentrations of NR-7h. Western blotting was used to evaluate the levels of p38-MAPK expression with untreated erythrocytes serving as the control and GAPDH as the loading control. **(ii)** Imaging flow cytometry was used to evaluate the fluorescence levels of p38-MAPK in the NR-7h (2.5µM) treated erythrocytes. Untreated erythrocytes were used as the control. **(B)** Erythrocytes were treated with NR-7h (2.5µM) in the presence and absence of MG132 proteasome inhibitor. Western blotting was used to evaluate p38-MAPK expression with GAPDH as the loading control. **(C) (i)** Uninfected and infected erythrocytes were treated with NR-7h (2.5µM) and the levels of p38MAPK expression were evaluated by Western blotting. Untreated erythrocytes were used as control and GAPDH as the loading control. **(ii)** *Pf*3D7 infected erythrocytes (at ring stage) were treated with different concentrations of NR-7h for 72hr. **(a)** Growth inhibition curve - %inhibition against log concentration of NR-7h. **(b)** Giemsa image panel of *P. falciparum*-infected erythrocytes (Scale bar: 10µm). **(iii)** Erythrocytes were pre-treated with 1µM and 5µM NR-7h and then infected with purified *Pf*3D7 schizonts. **(a)** Parasitemia plotted as % rings plotted against each NR-7h concentration. **(b)** Giemsa image panel of *P. falciparum*-infected erythrocytes (Scale bar: 5µm). Untreated infected erythrocytes served as the control group. Band intensities were analyzed using ImageJ software and plotted using GraphPad Prism 8. Statistical analysis utilized the unpaired t-test with Welch’s correction, with significance levels denoted as *, **, and *** for p-values less than 0.05, 0.01, and 0.001, respectively.

To determine the *in-vitro* antimalarial potency of NR-7h, *Pf*3D7 culture at the ring stage was treated with a range of NR-7h concentrations for 72 hours. Post-treatment, parasitemia was calculated for each group by Giemsa staining. NR-7h showed a concentration-dependent anti-malarial effect on *Pf*3D7 culture and the half-maximal inhibitory concentration (IC_50_) of NR-7h was determined to be 1.047 µM by the growth inhibition curve (Fig. 6C (ii)). Further, to determine the effect of NR-7h mediated p38-MAPK degradation on parasite invasion the NR-7h pre-treated erythrocytes were infected with purified *Pf*3D7 schizonts and the formation of rings was observed after 6-8 hours by Giemsa staining. A significant reduction was observed in the *Plasmodium* parasite’s ability to invade the NR-7h-treated erythrocytes at both 1µM (∼1.5-fold) and 5µM (∼ 3-fold) as compared to the untreated erythrocytes (Fig. 6C (iii)).

## Discussion

p38 MAPK is one of the critical members of the family of MAPKs and comprises four isoforms: p38α, p38β, p38γ, and p38δ, which differ in tissue expression and functional roles^50^. It is regulated by dual phosphorylation on its Thr-Gly-Tyr motif by upstream kinases that include MKK3 and MKK6 in response to stressors such as UV light, cytokines (eg, TNF-α, IL-1β), and PAMPs^51^. It influences many signaling pathways including inflammatory response, stress, apoptosis, differentiation, and transcription factor-mediated via ATF2, NF-κB, and proteins with the help of post-translational modification^52–55^. p38 MAPK has a dual role in cancer as a tumor suppressor and promoter^56^. It has been linked to chronic inflammatory diseases^57^, including rheumatoid arthritis and psoriasis^58^, and neurodegenerative disorders^59^, including Alzheimer’s^60^. Despite its value as a therapeutic target, clinical development of p38 MAPK inhibitors has been slow, limited by such factors as off-target effects and toxicity^57^.

In the present study, we observed the higher activation of human p38-MAPK at 6hr of *L. donovani* infection in human macrophages that further downregulates at later time points of infection suggesting the involvement of human p38-MAPK activation at the time of parasite entry. Using PROTAC-based approach, we degraded human p38-MAPK by NR-7h followed by *L. donovani* infection. We observed reduction of infected macrophages as well as the reduced number of parasites in those infected macrophages that indicates the requirement of human p38-MAPK by *L. donovani* not only for its invasion but also for its disease progression. p38-MAPK is a classical regulator of various cellular and immune mechanisms. For a better understanding of human p38-MAPK mediated host responses during *L. donovani* infection, we further investigated the effect of p38-MAPK degradation on host immune mechanisms in *L. donovani* infected macrophages.

Human p38-MAPK regulates cytokine production as its downstream process^49^, we observed that degradation of human p38-MAPK led to significant reduction in the levels of TNFα and IL-10, a pro-inflammatory and an anti-inflammatory cytokine, respectively. In the context of the *Leishmania* parasite infection, it is well-documented that parasites suppress host pro-inflammatory cytokine production^61,62^ and induce host anti-inflammatory cytokine production^63,64^ to facilitate their survival. Interestingly, when *Leishmania* parasites infected NR-7h-pre-treated macrophages, a significant upregulation of TNFα and a concomitant downregulation of IL-10 was observed. The data suggests the exploitation of host signaling in a p38-MAPK-mediated manner by the *L. donovani* parasites. Additionally, p38-MAPK also regulates oxidative stress^65^, therefore, we next observed ROS and NO levels, the two key processes to control the parasite. Degradation of human p38-MAPK induced generation levels of ROS and NO during *L. donovani* infection that might contribute in the reduction of parasite load. Since targeting human p38-MAPK is a host-directed therapeutic approach, we further evaluated the synergistic effect of NR-7h with the known antileishmanial drug Amphotericin B that acts directly on *Leishmania* parasites^66^. The combination of both compounds led to the near-complete clearance of *L. donovani* parasites.

In context to *Plasmodium falciparum* infection in human erythrocytes, a substantial reduction in parasitic load was observed in NR-7h treated erythrocytes with degraded p38-MAPK. During its intraerythrocytic stages of development, the *Plasmodium* parasite introduces osmotic changes by altering the permeability of the erythrocyte membrane^67^ and oxidative stress^68^ in the host erythrocytes, which serve as triggers for activation and downstream signaling of p38-MAPK. Activation of p38-MAPK under oxidative stress results in downstream phosphorylation of Src tyrosine kinase and Band 3 ultimately leading to lysis of erythrocytes^69^. Since the p38-MAPK activation is linked to erythrocyte lysis its inhibition is correlated to reduced lysis and parasite egress. Additionally, human erythrocytes with mutant Band 3, an invasion ligand of *Plasmodium* parasite^70^ lacking N-terminal 11 amino acids which otherwise carry the phosphorylation site show inhibited parasite invasion and maturation^71^. This further establishes the importance of the activation and signaling of p38-MAPK in infected erythrocytes. Previous studies have also shown that human p38-MAPK inhibitors including RWJ67657 and RWJ68198 affect the parasite development within the human erythrocytes^72^. Altogether, p38-MAPK degradation-mediated inhibition in parasite egress, invasion, and survival contributes to the overall reduction in parasitic load.

To conclude, this study establishes that NR-7h mediated degradation of host p38-MAPK significantly reduces *Leishmania donovani* and *Plasmodium falciparum* infections. p38-MAPK degradation modulates immune responses that effectively promotes parasite clearance (Fig. 7). Furthermore, host p38-MAPK degradation synergizes with Amphotericin B to effectively improve the clearance of *L. donovani* parasites, highlighting a promising therapeutic approach that could improve treatment outcomes for parasitic diseases.

**Fig. 7:**
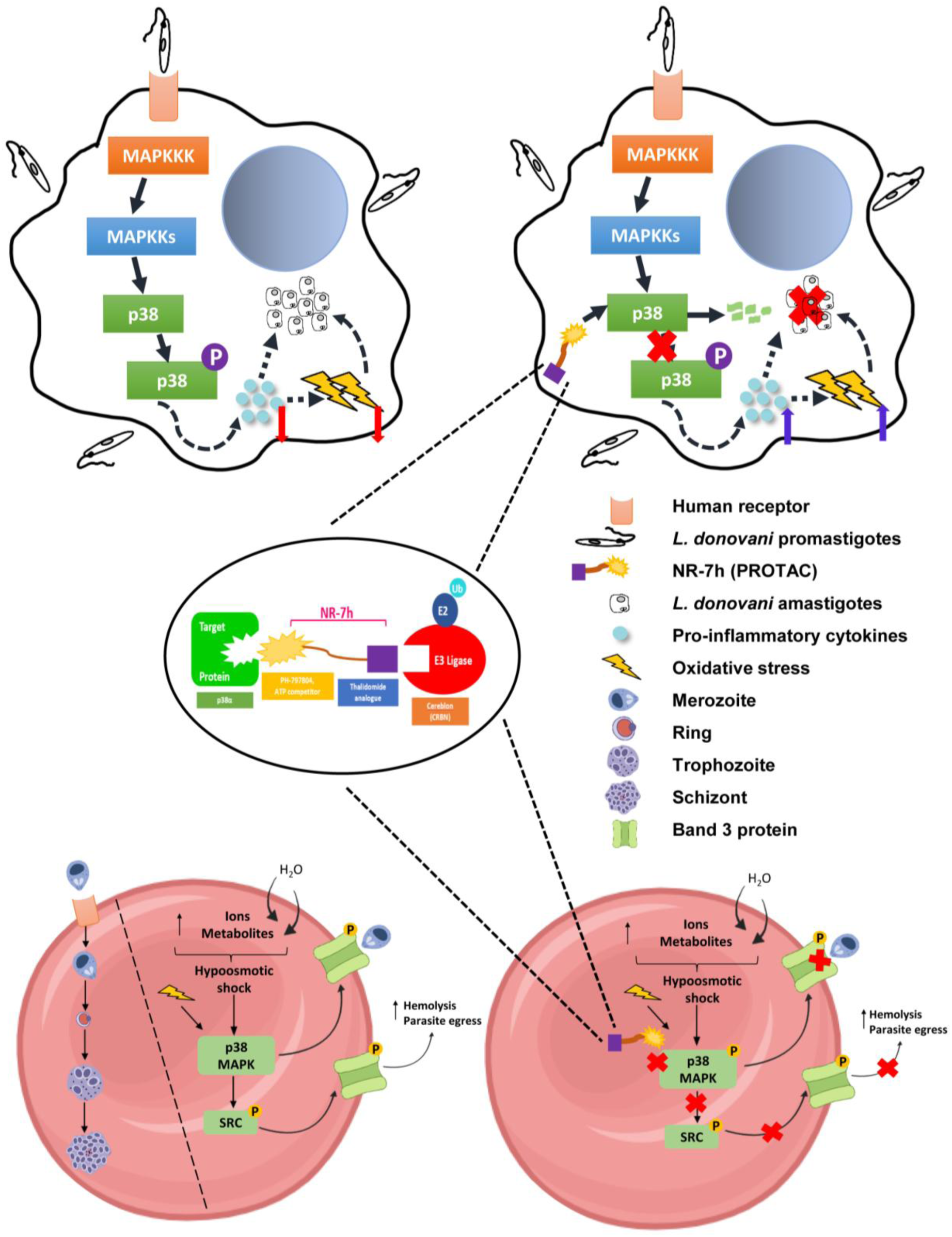
A schematic representation on the effect of host p38-MAPK degradation by NR-7h on the pathogenesis of *Leishmania donovani* and *Plasmodium falciparum* in macrophages and erythrocytes, their respective hosts.

## Materials and Methods

### Cell and parasite culture

THP-1 cells, obtained from the National Centre for Cell Science (NCCS, Pune, India), were cultured in RPMI-1640 media (Gibco) supplemented with 10% heat-inactivated fetal bovine serum (FBS) (Gibco), 2 mM L-glutamine, 10 mM HEPES buffer, 20 mM sodium bicarbonate, 1 mM sodium pyruvate, and penicillin/streptomycin (10,000 units/ml) at 37°C in a humidified incubator with 5% CO_2_. To induce differentiation into macrophages, Phorbol 12-myristate 13-acetate (PMA) (Sigma) was added at a concentration of 50 ng/mL for 24hr. *L. donovani* Bob promastigotes were cultured in M199 medium (Gibco) supplemented with 10% FBS (Gibco) and 10 µg/mL gentamycin at 26°C. Infections were performed using metacyclic stage of *L. donovani* promastigotes at a multiplicity of infection (MOI) of 20.

### Protein immunoblotting

To prepare cell lysates, RIPA lysis buffer (GBiosciences) supplemented with a protease inhibitor cocktail (Roche) was used. Equal amounts of cell lysates (25µg for THP-1 macrophages and 100µg for erythrocytes) were loaded and separated using SDS-PAGE. The proteins were then transferred onto a nitrocellulose immunoblot membrane (Bio-Rad). The membrane was blocked overnight at 4°C in 5% BSA (SRL) in 1X PBS containing 0.05% Tween20 (Sigma) (wash buffer), followed by three washes with the wash buffer. Subsequently, the membrane was incubated with primary antibodies for 2hr at room temperature, followed by three additional washes and incubation with HRP-conjugated secondary antibodies for another 1hr at room temperature. After three washes, the blots were visualized using a densitometric ECL kit (Bio-Rad) on the ChemiDoc Imaging System (Bio-Rad). The data was analyzed using ImageJ software.

### Imaging Flow Cytometry

THP-1 macrophages were treated with 1μM NR-7h along the co-stimulation of LPS (100ng/mL) for 6hr. Cells were fixed, permeabilized, and then stained with 1:200 of each human p38-MAPK (Biolegend #622403) and human Phospho-p38-MAPK (Biolegend #690201) for an hour followed by the secondary antibodies, anti-rabbit Alexa fluor 488 (Invitrogen #A11008) and anti-mouse Alexa fluor 546 (Invitrogen #A11030), respectively for half an hour. Similarly, erythrocytes were treated with 2.5µM NR-7h to analyze the p38-MAPK degradation by imaging flow cytometry. The cells were fixed in cold 2% paraformaldehyde and permeabilized in 0.001% Triton X-100. Subsequently, the cells were stained with human p38-MAPK antibody (1:200) for 1.5hr and anti-mouse Alexa 488 antibody (1:200) for 45 minutes. After washing, the cells were processed in Cytek Amnis FlowSight imaging flow cytometer and the data was analyzed using Ideas Software.

### Cell viability assessment

MTT assay was conducted to evaluate the impact of NR-7h on the viability of THP-1 cells. A solution of MTT dye (Sigma-Aldrich) was prepared by diluting 5 mg of MTT in 1 mL of PBS, which was further diluted to 1:10 in RPMI medium. For the assay, 10,000 THP-1 cells in 100µL of complete media were seeded in each well of 96-well flat-bottom plates. Following the differentiation of monocytes into macrophages, THP-1 macrophages were treated with various concentrations of NR-7h with the co-stimulation of LPS (100ng/mL). After 72hr, the MTT assay was carried out according to the manufacturer’s instructions. Specifically, treated or untreated THP-1 macrophages were incubated with MTT dye solution for 2hr at 37°C, followed by adding a stopping buffer (5% formic acid in isopropanol) to halt the reaction for 20 minutes at 37°C. Absorbance was then measured at 570 nm, and the cell viability percentage was calculated. Each experiment was performed in triplicates and repeated three times.

### RNA Extraction and Gene Expression Profiling

Isolation of total RNA from treated or untreated macrophages was performed at an appropriate time point using the TRIZOL reagent (Invitrogen). Subsequently, the quantification of the isolated RNA was conducted using a Nanodrop ND-1000 spectrophotometer (Thermo Fisher Scientific). For cDNA synthesis, one microgram of total RNA was utilized and the process was carried out using the First Strand cDNA Synthesis Kit (Thermo Fisher Scientific) according to the manufacturer’s instructions, employing random hexamer primers. The PCR reactions were performed on an Applied Biosystems Real-Time PCR System (ABI) using the PowerUp SYBR Green PCR Master Mix (Thermo Fisher Scientific).

### Parasite quantification via Propidium iodide staining

0.5X10^6^ THP-1 cells were cultured on 18 mm diameter round coverslips coated with Poly-L-Lysine (Sigma) and treated with 50 ng/mL PMA to induce macrophage differentiation for 24hr. Following the incubation, the coverslips were rinsed with RPMI-1640 medium to eliminate non-adherent cells. Subsequently, the adherent THP-1 macrophages were treated with different concentrations of NR-7h with the co-stimulation of LPS (100ng/mL) for 6hr, followed by infection with metacyclic *L. donovani* promastigotes at a multiplicity of infection (MOI) of 20. After 6hr of infection, the cells were washed three times with RPMI medium to remove non-phagocytosed promastigotes. The infected macrophages were then further incubated for 24hr. Following the incubation, the coverslips were washed with PBS and fixed with chilled methanol. Subsequently, the cells were stained with propidium iodide (500 nM in 2× SSC buffer)^16^. A minimum of 100 macrophages per coverslip were counted using a confocal microscope to determine the number of resident amastigotes. Confocal imaging was performed using a High-speed laser scanning confocal microscope with a 60x objective magnification, utilizing an excitation/emission wavelength of 535/617 nm.

### Immunofluorescence assay

THP-1 cells were seeded on 18-mm-diameter round coverslips coated with poly-L-lysine (Sigma) and were stimulated with 50 ng/mL of PMA to differentiate into macrophages for 24hr. THP-1 macrophages were treated with 1μM NR-7h with the co-stimulation of LPS (100ng/mL) for 6hr, followed by infection with metacyclic *L. donovani* promastigotes at a MOI of 20 for 12hr, 36hr, and 60hr. Cells were washed with RPMI followed by fixation with 2% paraformaldehyde (PFA) for 15 min. Cells were washed with 1× PBS and were permeabilized with a permeabilization buffer (0.1% BSA and 0.2% saponin in 1× PBS). Then, cells were washed with washing buffer (0.1% BSA and 0.1% saponin in 1× PBS) followed by the incubation of 1:200 dilution of primary antibody for p38-MAPK at room temperature for 1hr. Following washes, a secondary antibody, anti-rabbit Alexa Fluor 488 at a dilution of 1:250 was introduced to the cells for 30 min at room temperature. Cells were washed and stained with propidium iodide (500 nM in 2× SSC buffer). The coverslips were mounted on the slides with a mounting medium, fluoroshield with DAPI (Sigma). The fluorescence intensity of p38-MAPK and the number of resident amastigotes were determined using confocal microscopy.

### Quantification of Reactive Oxygen Species (ROS)

THP-1 differentiated human macrophages were treated with 1μM NR-7h along the co-stimulation of LPS (100ng/mL) for a duration of 6hr in 96 well plates (Corning #CLS3603). Following the treatment, macrophages were infected with *L. donovani* promastigotes at a multiplicity of infection (MOI) of 20 for 30 minutes at 37°C. Simultaneously, cells were loaded with 10µM DCFH-DA for 30 minutes, following the guidelines provided by the manufacturer (Abcam #ab113851). Once the incubation period concluded, the cells were promptly analyzed for levels of ROS generation using fluorimetry with an excitation/emission of 485 nm/535 nm.

### Nitric Oxide (NO) quantification

PMA differentiated THP-1 macrophages were treated with 1μM NR-7h along the co-stimulation of LPS (100ng/mL) for a duration of 6hr, followed by infection with *L. donovani* promastigotes at an MOI of 20 along the co-stimulation with LPS (100ng/mL) and hIFNγ (20ng/mL). After 24hr post-infection, the supernatant was collected to determine the level of NO (in the form of nitrite) using the Griess reagent kit (Invitrogen #G-7921), following the instructions provided by the manufacturer.

### *Plasmodium* parasite culture and *in-vitro* assays

*Plasmodium falciparum* 3D7 (*Pf*3D7) was cultured in O+ human erythrocytes and RPMI 1640 (Gibco) supplemented with 0.5% albumax (Gibco), 2g/L sodium bicarbonate (Biobasic), 30mg/L hypoxanthine (Sigma) and 10mg/L gentamycin (SRL) under mixed gas conditions. *In-vitro* growth inhibition assay was carried out in synchronized *Pf*3D7 culture at ring stage at 0.8% parasitemia and 2% hematocrit. *Pf*-infected erythrocytes were treated with different concentrations of NR-7h (20µM, 10µM, 5µM, 2.5µM, 1.25µM, 625nM, 312nM, and 156nM) for 72hr in a total volume of 100µL in each well of 96-well microtiter plate. Post 72hr, Giemsa smears were prepared, and parasitemia for each group was calculated by counting at least 2000 cells. Growth inhibition ability of NR-7h was calculated as % inhibition and it was plotted against logarithmic NR-7h concentration. For *in-vitro Pf*3D7 invasion assay, erythrocytes were pre-treated with 1µM and 5µM. Post-incubation with NR-7h, the erythrocytes were washed with 1X iRPMI to remove the residual drug. Purified *Pf*3D7 schizonts were incubated to pre-treated erythrocytes at 0.8% parasitemia and 1% hematocrit in a total volume of 100µL in each well of a 96-well microtiter plate. 6-8hr post-incubation, Giemsa smears were prepared for each group. Formation of ring in the infected erythrocyte was observed and parasitemia was plotted as % rings against each NR-7h concentration. Untreated infected erythrocytes served as the control group in the *in-vitro* assays. For the in-vitro assays, parasitemia was calculated by the formula: *% Parasitemia = (Infected erythrocytes / Total number of erythrocytes) X 100,* while the % inhibition was calculated as *% inhibition = [(Parasitemia_control_ - Parasitemia_Treated_)] / Parasitemia_control_ X 100*

### Data Analysis and Interpretation

The data were displayed as the mean±SD (standard deviation). Every experiment was replicated thrice in distinct batches. All charts produced and the corresponding statistical evaluations were carried out utilizing GraphPad Prism (GraphPad Software, USA). The statistical significance was determined using the unpaired t-test with Welch’s correction. The significance level was achieved with p-values <0.05. The p-values were denoted as *p < 0.05, **p < 0.01, ***p < 0.001, and ****p < 0.0001.

## Data availability

The original contributions presented in the study are included in the article. Further inquiries can be directed to the corresponding authors.

## Acknowledgements

We thank Prof. Angel R. Nebreda and Prof. Antoni Riera from the Institute for Research in Biomedicine, Barcelona for generously providing NR-7h, a PROTAC for p38-MAPK. We also thank the National Center for Cell Science, Pune, for providing THP-1 cells. We are grateful to JNU for providing access to the anti-plagiarism software Turnitin, and also thank the Advanced Instrumentation Research Facility, JNU, for providing access to a confocal microscopy facility. Cytek Amnis FlowSight imaging flow cytometer used in the study was supported by DBT-Indo-Swiss sanctioned to S.S. (IC-12044(11)/10/2021-ICD-DBT). N.R., R.B., and P.S. are supported by DBT-JRF, DHR-Women Scientist, and ICMR-SRF respectively. J.S. is a recipient of the BioCARe Women Scientist fellowship from DBT. S.S. is a recipient of the National Bio Scientist Award. The Science and Engineering Research Board has funded this work under the Department of Science and Technology (IPA/2020/000007) sanctioned to A.R. and S.S.

## Author contributions

A.R. and S.S. conceptualized the idea, designed the experiments, executed the data interpretation, and analyzed the experimental data. A.R. and S.S. reviewed the work and corrected the manuscript. J.S., N.R., and S.S. wrote the manuscript. J.S. and N.R. designed the study, conducted the investigation, curated, analyzed and interpreted the data. J.S. performed degradation and immune modulation studies in macrophages. N.R. performed degradation and anti-malarial studies in RBC. R.B. and J.S. performed invasion studies. P.S. and G.D.G.P. helped in the *Leishmania-*related experimental methodology. R.P.P. assisted in the interpretation of *Leishmania*-related experimental data. J.S., N.R., A.R., and S.S. critically analyzed the data. All authors contributed to the article and approved the submitted version.

## Competing interests

The authors declare no competing interests.

## Supplementary information

## Notes

### Competing Interest Statement

The authors have declared no competing interest.

